# Genetic architecture of early childhood growth phenotypes gives insights into their link with later obesity

**DOI:** 10.1101/150516

**Authors:** N. Maneka G. De Silva, Sylvain Sebert, Alexessander Couto Alves, Ulla Sovio, Shikta Das, Rob Taal, Nicole M. Warrington, Alexandra M. Lewin, Marika Kaakinen, Diana Cousminer, Elisabeth Thiering, Nicholas J. Timpson, Ville Karhunen, Tom Bond, Xavier Estivill, Virpi Lindi, Jonathan P. Bradfield, Frank Geller, Lachlan J.M. Coin, Marie Loh, Sheila J. Barton, Lawrence J. Beilin, Hans Bisgaard, Klaus Bønnelykke, Rohia Alili, Ida J. Hatoum, Katharina Schramm, Rufus Cartwright, Marie-Aline Charles, Vincenzo Salerno, Karine Clément, Cornelia M. van Duijn, Elena Moltchanova, Johan G. Eriksson, Cathy Elks, Bjarke Feenstra, Claudia Flexeder, Stephen Franks, Timothy M. Frayling, Rachel M. Freathy, Paul Elliott, Elisabeth Widén, Hakon Hakonarson, Andrew T. Hattersley, Alina Rodriguez, Marco Banterle, Joachim Heinrich, Barbara Heude, John W. Holloway, Albert Hofman, Elina Hyppönen, Hazel Inskip, Lee M. Kaplan, Asa K. Hedman, Esa Läärä, Holger Prokisch, Harald Grallert, Timo A. Lakka, Debbie A. Lawlor, Mads Melbye, Tarunveer S. Ahluwalia, Marcella Marinelli, Iona Y. Millwood, Lyle J. Palmer, Craig E. Pennell, John R. Perry, Susan M. Ring, Markku Savolainen, Kari Stefansson, Gudmar Thorleifsson, Fernando Rivadeneira, Marie Standl, Jordi Sunyer, Carla M.T. Tiesler, Andre G. Uitterlinden, Inga Prokopenko, Karl-Heinz Herzig, George Davey Smith, Paul O'Reilly, Janine F. Felix, Jessica L. Buxton, Alexandra I.F. Blakemore, Ken K. Ong, Struan F.A. Grant, Vincent W.V. Jaddoe, Mark I. McCarthy, Marjo-Riitta Järvelin

**Affiliations:** Department of Epidemiology and Biostatistics, MRC-PHE Centre for Environment and Health, School of Public Health, Imperial College London, UK.; Center for Life Course Health Research, Faculty of Medicine, University of Oulu 90014 Oulu, Finland.; Biocenter Oulu, University of Oulu, Finland.; Department of Obstetrics and Gynaecology, University of Cambridge, Cambridge, UK.; Department of Paediatrics, Erasmus Medical Center, Sophia Children's Hospital, Rotterdam, Netherlands.; The Generation R Study Group, Erasmus Medical Center, Rotterdam, Netherlands.; School of Women’s and Infants’ Health, The University of Western Australia, Australia.; The University of Queensland Diamantina Institute, The University of Queensland, Translational Research Institute, Brisbane, Queensland, 4102, Australia.; Department of Mathematics, Brunel University, London, UK.; Department of Genomics of Common Disease, School of Public Health, Imperial College London, Hammersmith Hospital, London, UK.; Centre for Pharmacology and Therapeutics, Division of Experimental Medicine, Department of Medicine, Imperial College London, Hammersmith Hospital, London, UK; Division of Human Genetics, The Children's Hospital of Philadelphia, Philadelphia, Pennsylvania, USA.; Institute of Epidemiology I, Helmholtz Zentrum München -, German Research Center for Environmental Health, Munich Neuherberg, Germany.; Ludwig-Maximilians-University of Munich, Dr. von Hauner Children's Hospital, Division of Metabolic Diseases and Nutritional Medicine, Dr von Hauner Children's Hospital, Ludwig-Maximilians University Munich, Munich, Germany.; MRC Integrative Epidemiology Unit at the University of Bristol, UK.; School of Social and Community Medicine, University of Bristol.; Genomics and Disease Group, Bioinformatics and Genomics Programme, Centre for Genomic Regulation (CRG), Barcelona, Catalonia, Spain.; Pompeu Fabra University (UPF), Barcelona, Catalonia, Spain.; Hospital del Mar Medical Research Institute (IMIM), Barcelona, Catalonia, Spain.; Spanish consortium for Research on Epidemiology and Public Health (CIBERESP), Spain.; Sidra Medical and Research Center, Doha, Qatar.; Institute of Biomedicine, Department of Physiology, University of Eastern Finland, Kuopio, Finland.; Center for Applied Genomics, Abramson Research Center, The Children's Hospital of Philadelphia, Philadelphia, Pennsylvania, USA.; Department of Epidemiology Research, Statens Serum Institut, Copenhagen, Denmark.; Institute for Molecular Bioscience, The University of Queensland, Brisbane, Australia.; Translational Laboratory in Genetic Medicine (TLGM), Agency for Science, Technologyand Research (A*STAR), 8A Biomedical Grove, Immunos, Level 5, Singapore 138648.; MRC Lifecourse Epidemiology Unit, University of Southampton, Southampton General Hospital, Southampton, UK.; School of Medicine and Pharmacology, Royal Perth Hospital, The University of Western Australia, Australia.; COPSAC, The Copenhagen Prospective Studies on Asthma in Childhood, Faculty of Health Sciences, University of Copenhagen, Copenhagen, Denmark.; The Danish Pediatric Asthma Center, Copenhagen University Hospital, Gentofte, Denmark.; CRNH Ile de France, Hôpital Pitié-Salpêtrière, Paris, France.; Obesity, Metabolism, and Nutrition Institute and Gastrointestinal Unit, Massachusetts General Hospital, Boston, MA, USA.; Department of Medicine, Harvard Medical School, Boston, MA, USA.; Institute of Human Genetics, Helmholtz Center Munich, German Research Center for Environmental Health, Neuherberg, Germany.; Institute of Human Genetics, Technische Universität München, München, Germany.; Institute for Reproductive and Developmental Biology, Imperial College London, UK.; Inserm, UMR 1153 (CRESS), Villejuif; Paris Descartes University, France.; Department of Epidemiology, Erasmus Medical Center, Rotterdam, Netherlands.; University of Canterbury, Department of Mathematics and Statistics, Christchurch, New Zealand.; Department of General Practice and Primary Health Care, University of Helsinki, and Helsinki University Hospital, Helsinki, Finland.; Department of Chronic Disease Prevention, National Institute for Health and Welfare, Helsinki, Finland.; Folkhalsan Research Center, Helsinki, Finland.; MRC Epidemiology Unit, University of Cambridge School of Clinical Medicine, Box 285 57, Institute of Metabolic Science, Cambridge Biomedical Campus, Cambridge, CB2 0QQ, UK.; Institute of Biomedical and Clinical Science, University of Exeter Medical School, University of Exeter, Royal Devon and Exeter Hospital, Exeter EX2 5DW, UK.; Institute for Molecular Medicine Finland, University of Helsinki, Helsinki, Finland.; Department of Paediatrics, Perelman School of Medicine, University of Pennsylvania, Philadelphia, Pennsylvania, USA.; Institute of Diabetes, Obesity and Metabolism, Perelman School of Medicine, University of Pennsylvania, Philadelphia, Pennsylvania, USA.; School of Psychology, College of Social Science, University of Lincoln Brayford Pool Lincoln, Lincolnshire, LN6 7TS, UK.; Human Genetics and Medical Genomics, Faculty of Medicine, University of Southampton, UK.; School of Population Health, University of South Australia, Adelaide, Australia.; Centre for Paediatric Epidemiology and Biostatistics, University College London Institute of Child Health, London, UK.; Wellcome Trust Centre for Human Genetics, University of Oxford, UK.; Cardiovascular Medicine unit, Department of Medicine, Karolinska Institute, Stockholm, Sweden.; Research Unit of Mathematical Sciences, University of Oulu, Finland.; Research Unit of Molecular Epidemiology, Helmholtz Zentrum München, German Research Center for Environmental Health, Neuherberg, Germany.; German Center for Diabetes Research (DZD). Neuherberg, Germany.; Kuopio Research Institute of Exercise Medicine, Kuopio, Finland.; Department of Clinical Physiology and Nuclear Medicine, Kuopio University Hospital, Kuopio, Finland.; ISGlobal, Centre for Research in Environmental Epidemiology (CREAL), Barcelona, Spain Center for Research in Environmental Epidemiology (CREAL), Barcelona, Catalonia, Spain.; Clinical Trial Service Unit and Epidemiological Studies Unit (CTSU), University of Oxford, Old Road Campus, UK.; Medical Research Council Population Health Research Unit (MRC PHRU) at the University of Oxford, Oxford, UK.; School of Public Health and Robinson Research Institute, University of Adelaide, Australia.; Avon Longitudinal Study of Parents and Children, School of Social and Community Medicine, University of Bristol, UK.; Division of Internal Medicine, and Biocenter of Oulu, Faculty of Medicine, Oulu University, Finland.; deCODE genetics, Reykjavik, Iceland.; University of Iceland, Faculty of Medicine, Iceland.; Department of Internal Medicine, Erasmus Medical Center, Rotterdam, Netherlands.; Oxford Centre for Diabetes, Endocrinology and Metabolism, University of Oxford, Churchill Hospital, Old Road, Headington, Oxford, UK.; Research Unit of Biomedicine, and Biocenter of Oulu, Oulu University, 90014 Oulu, Finland.; Medical Research Center and Oulu University Hospital, University of Oulu and Oulu University Hospital, Oulu, Finland.; Department of Gastroenterology and Metabolism, Poznan University of Medical Sciences, Poznan, Poland.; MRC Social, Genetic and Developmental Psychiatry Centre, Institute of Psychiatry, King’s College London, De Crespigny Park, London, UK.; UCL Genetics Institute, Department of Genetics, Evolution and Environment, University College London, London, UK.; Department of Life Sciences, College of Health and Life Sciences, Brunel University London, London, UK.; Oxford NIHR Biomedical Research Centre, Churchill Hospital, Old Road, Headington, Oxford, UK.; Unit of Primary Care, Oulu University Hospital OYS, Finland.; Department of Children and Young People and Families, National Institute for Health and Welfare, Oulu, Finland.

**Author notes:** These authors equally contributed to the work (N. Maneka G. De Silva, Sylvain Sebert, Alexessander Couto Alves, Ulla Sovio, Shikta Das, Rob Taal). These authors jointly directed this work (Struan F.A. Grant, Vincent W.V. Jaddoe, Mark I. McCarthy, Marjo-Riitta Järvelin). Correspondence should be addressed to M.-R.J., M.I.M., S.F.A.G., V.W.V.J.

## Abstract

Early childhood growth patterns are associated with adult metabolic health, but the underlying mechanisms are unclear. We performed genome-wide meta-analyses and follow-up in up to 22,769 European children for six early growth phenotypes derived from longitudinal data: peak height and weight velocities, age and body mass index (BMI) at adiposity peak (AP ^~^9 months) and rebound (AR ^~^5-6 years). We identified four associated loci (*P*< 5x10^−8^): *LEPR/LEPROT* with BMI at AP, *FTO* and *TFAP2B* with Age at AR and *GNPDA2* with BMI at AR. The observed AR-associated SNPs at *FTO, TFAP2B* and *GNPDA2* represent known adult BMI-associated variants. The common BMI at AP associated variant at *LEPR/LEPROT* was not associated with adult BMI but was associated with *LEPROT* gene expression levels, especially in subcutaneous fat (*P*<2x10^−51^). We identify strong positive genetic correlations between early growth and later adiposity traits, and analysis of the full discovery stage results for Age at AR revealed enrichment for insulin-like growth factor 1 (IGF-1) signaling and apolipoprotein pathways. This genome-wide association study suggests mechanistic links between early childhood growth and adiposity in later childhood and adulthood, highlighting these early growth phenotypes as potential targets for the prevention of obesity.

## Introduction

Complex developmental processes regulate changes in body mass and growth in early life. Multiple genetic variants are robustly associated with adult body mass index (BMI)^1^ and metabolic phenotypes ^2^^-^^4^ but much fewer data exist for childhood traits^5,6^. Discrete genetic variants may regulate developmental patterns during rapid growth in infancy compared to later growth periods in childhood^7^ as demonstrated by the age-dependent impact of variation at the *FTO* locus on BMI^8^ but there still remains a paucity of relevant data. On the other hand, robust observational associations between early growth traits and adult cardiometabolic risk ^9-12^ may, in part, be explained by shared genetic factors. In support of this hypothesis, studies from the Early Growth Genetics (EGG) consortium show that some of the genetic component contributing to size at birth also impacts type 2 diabetes, cardiovascular disease and blood pressure^13^. However, the extent of the genetic correlation between postnatal growth patterns and the regulation of adult body weight, energy metabolism and later metabolic health is unclear.

In infancy and childhood, individuals follow well-characterized and predictable height, weight and BMI trajectories (**Figure 1**)^14^. For example, BMI trajectory encompasses three periods characterized by i) a rapid increase in BMI up to the age of 9 months (adiposity peak, AP); ii) a rapid decline in BMI up to the age of 5-6 years (adiposity rebound, AR); and iii) a steady increase in BMI after 5-6 years until early adulthood^10^. Here, we set out to model sex-specific individual weight, height and BMI curves in children, using unique data collected from primary health care or clinical research visits, to extract six early growth traits: peak height velocity (PHV; cm/month), peak weight velocity (PWV; kg/month), age at adiposity peak (Age-AP; year), BMI at adiposity peak (BMI-AP, kg/m^2^), age at adiposity rebound (Age-AR; year), BMI at adiposity rebound (BMI-AR, kg/m^2^) (**Figure 1**) and to test for genetic associations using genome-wide genotype data. Although a few previous studies have investigated the genetic basis of these well-established measures of longitudinal growth trajectory^7,9,15,16^, no systematic genome-wide analyses have been carried out to date. Understanding the genetic architecture of early childhood growth phenotypes will provide insight into their regulation and into the mechanisms underlying the observed associations with adult cardiometabolic disease, which will be fundamental for early promotion of metabolic health.

**Figure 1.**
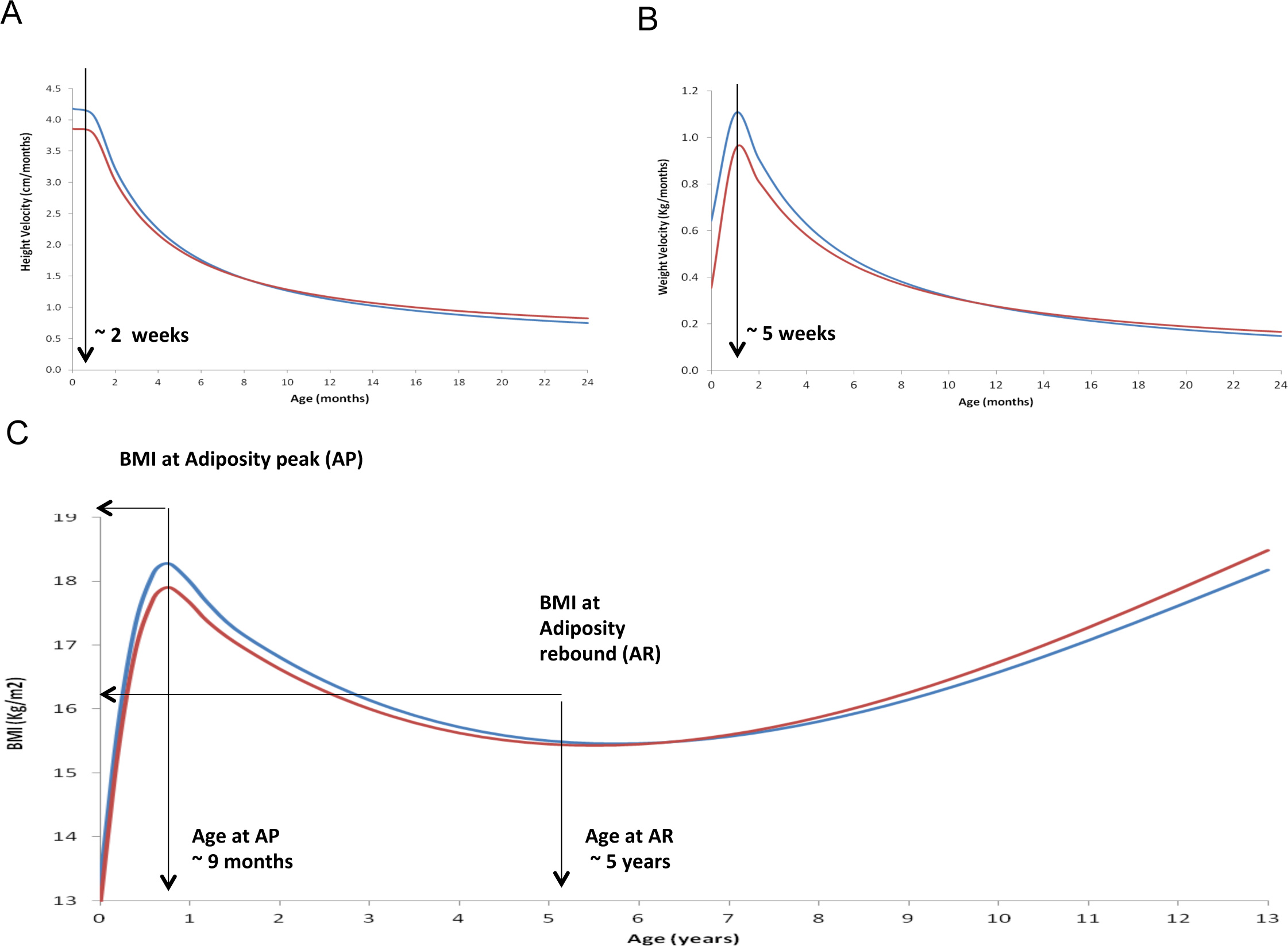
Graphical illustration of height and weight growth patterns and the derived measures of early growth traits used in the present study. A) Peak Height Velocity (PHV); B) Peak Weight Velocity (PWV); C) Age and BMI at Adiposity Peak (AP) and Rebound (AR). The growth curves for males are in blue and for females in red. Curves an based on the Northern Finland Birth Cohort (NFBC) 1986 data.

## Results

### Genome-wide association meta-analyses of early growth phenotypes

We conducted genome-wide association study (GWAS) meta-analyses to identify genetic loci influencing each of the six early growth traits in up to 7,215 term-born children of European ancestry from up to four population-based studies that had both genetic and early growth trait data (stage 1; **Online Methods**, **Supplementary Table 1, Supplementary Figure 1**). From the stage 1 inverse variance meta-analyses, we selected a total of 8 loci with either *P* < 1 x 10^−7^ or with *P* < 1 x 10^−5^ in/near genes known to be associated with obesity and metabolic traits in published GWAS or candidate gene studies (**Supplementary Table 2**), and sought confirmation in up to 16,550 term-born children from up to 11 additional studies (stage 2; **Online Methods, Supplementary Table 3, Supplementary Figure 1**). Our study design is summarized in **Figure 2**, while participant characteristics, genotyping arrays, imputation and quality control for the discovery and follow-up studies are summarized in **Supplementary Tables 1 and 3**. In combined meta-analyses of the discovery and follow-up studies (including up to 22,769 children), we identified common variants at four independent loci, each associated with any of the early growth phenotypes, at genome-wide levels of significance (i.e. *P* < 5 x 10^−8^) (Table 1, Figure 3, **Supplementary Figure 2**). The lead signals at each of the four loci were: rs9436303 at the *LEPR/LEPROT* (encoding the leptin receptor and an overlapping transcript) locus associated with BMI at AP, rs1421085 at the *FTO* (encoding a 2-oxoglutarate-dependent demethylase) and rs2817419 at the *TFAP2B* (encoding transcription factor AP-2 beta) loci associated with Age at AR, and rs10938397 near the *GNPDA2* (encoding adiposity regulating glucosamine-6-phosphate deaminase) locus associated with BMI at AR. The lead signals at each of the four loci were located within non-coding regions of the relevant genes except for rs10938397, which was located in an intergenic region proximal to the *GNPDA2* gene (**Figure 3**). The common AR-associated SNPs at *FTO, TFAP2B* and *GNPDA2* have previously been associated with other anthropometric measures, especially in adulthood^1^. However, the three loci *LEPR/LEPROT*, *TFAP2B* and *GNPDA2* described here are novel with respect to their association with early growth phenotypes, while the association between SNPs at *FTO* and Age at AR in the current study replicates the findings of a previous gene-centric study^8^ at genome-wide levels of significance.

**Table 1.**
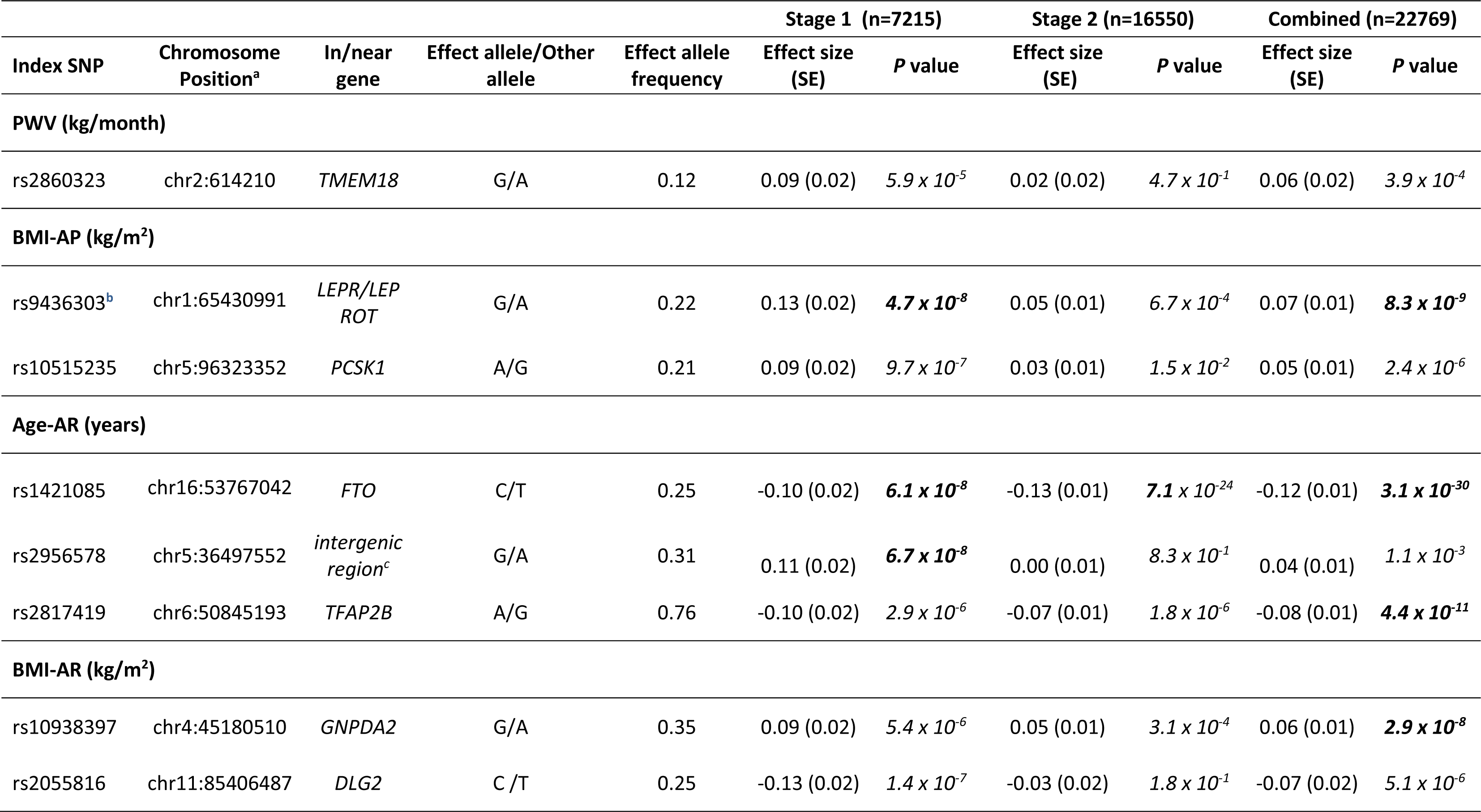
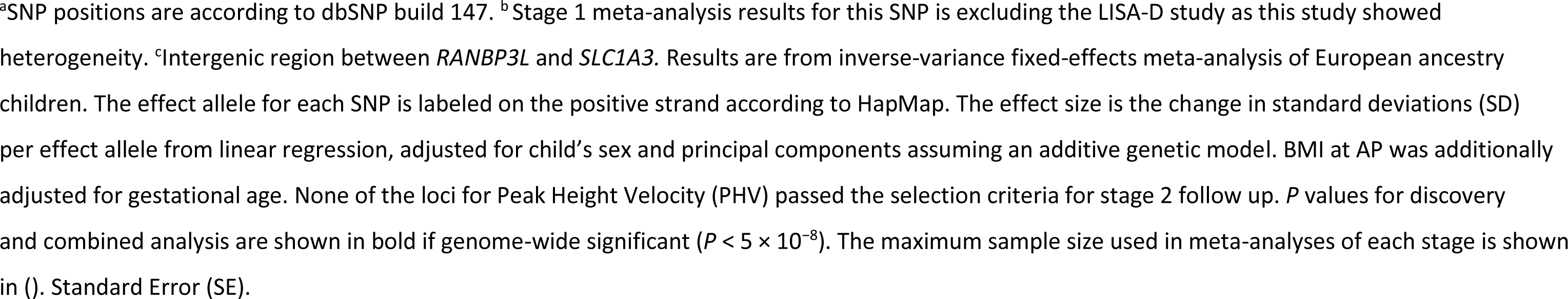
Summary statistics of the eight independent SNPs associated with Peak Weight Velocity in infancy (PWV), BMI at Adiposity Peak in infancy (BMIAP), Age at Adiposity Rebound (Age-AR) and BMI at Adiposity Rebound (BMI-AR) in discovery (stage 1), follow-up (stage 2) and in combined meta-analyses.

**Figure 2.**
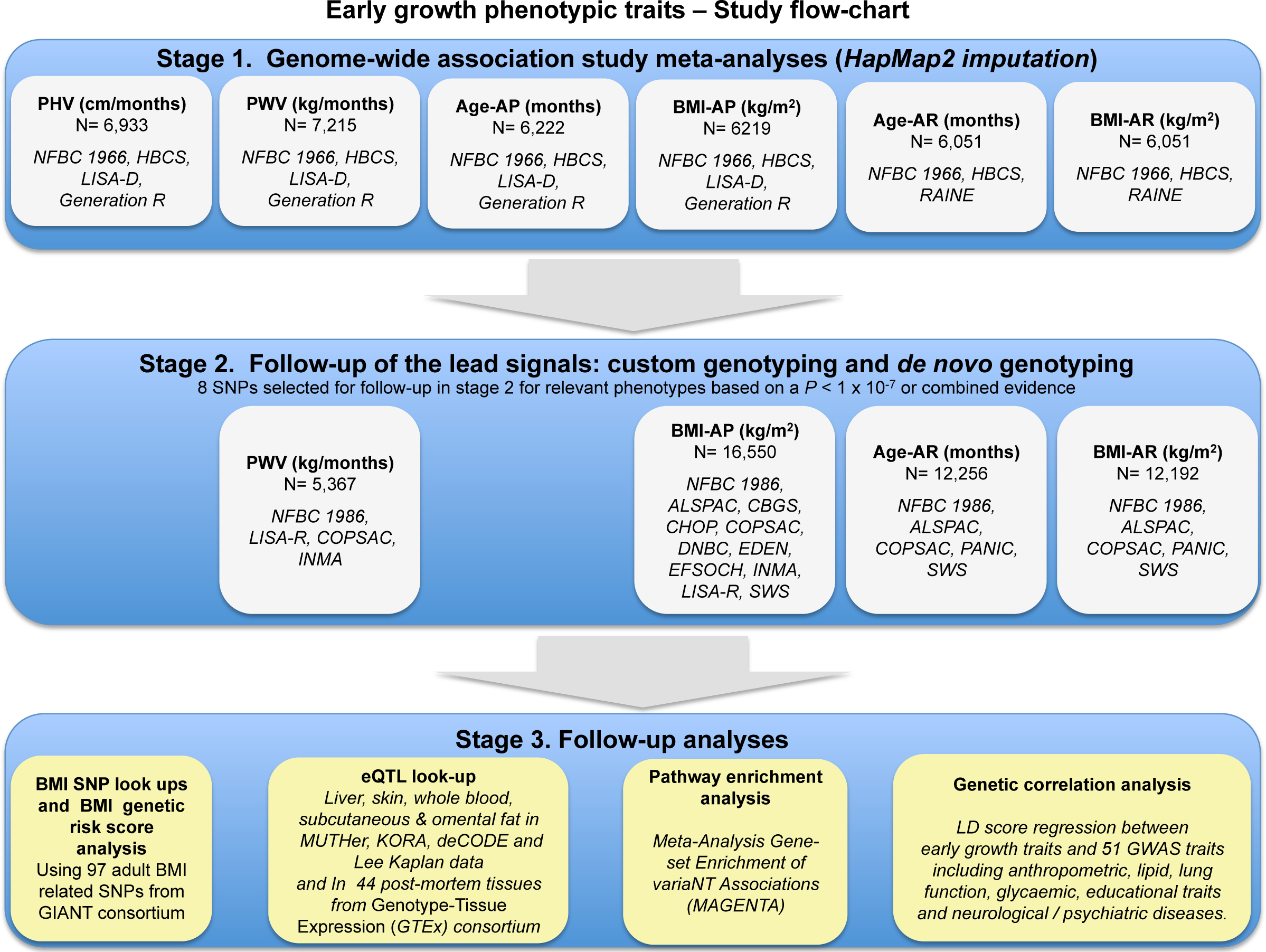
Summary of study design. The Flowchart shows the studies and total sample size used in Stage 1 genome-wide association meta-analyses and in Stage 2 follow u for each early growth trait leading to Stage 3 follow-up analyses. Stage 1 included up to 7215 European ancestry children from up to four population-based studies and stage 2 follow up includedup to 16,550 European ancestry children from up to 11 studies.

**Figure 3.**
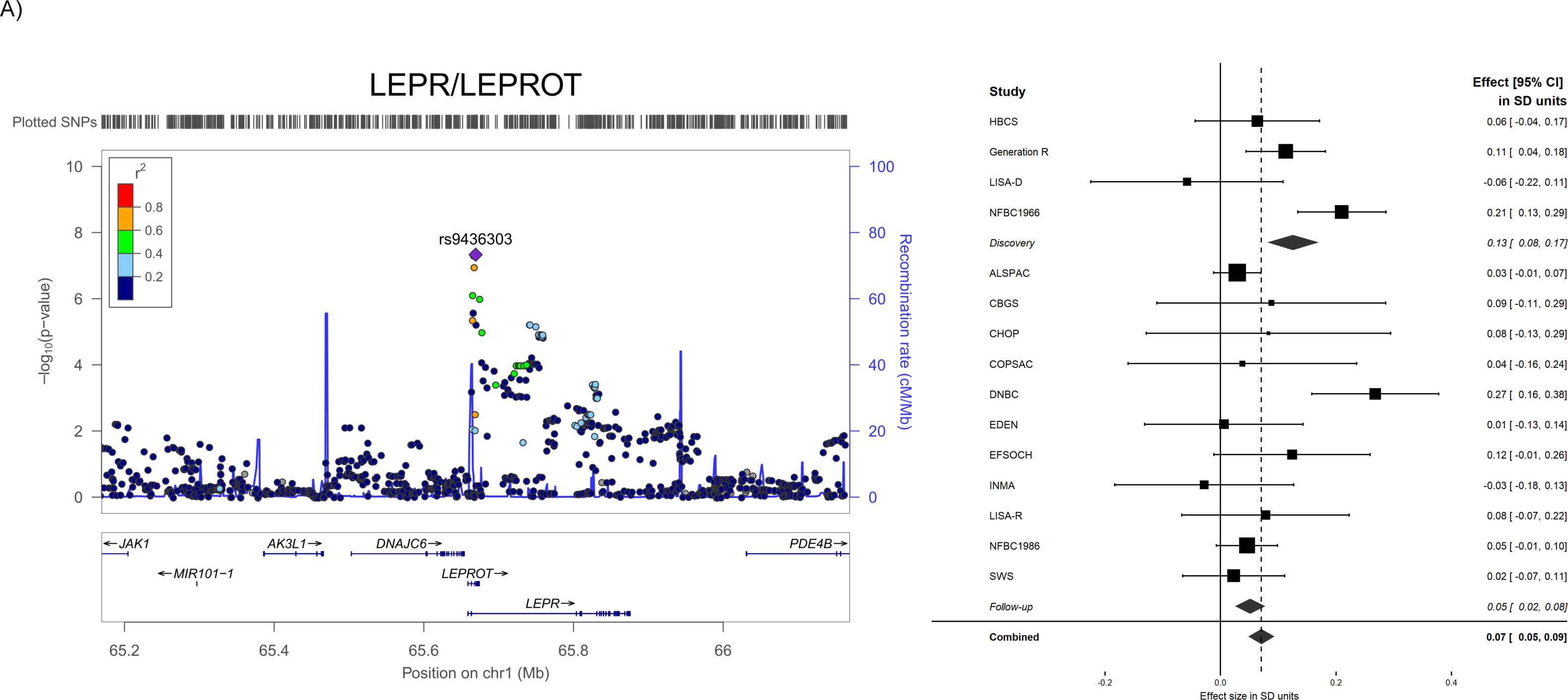
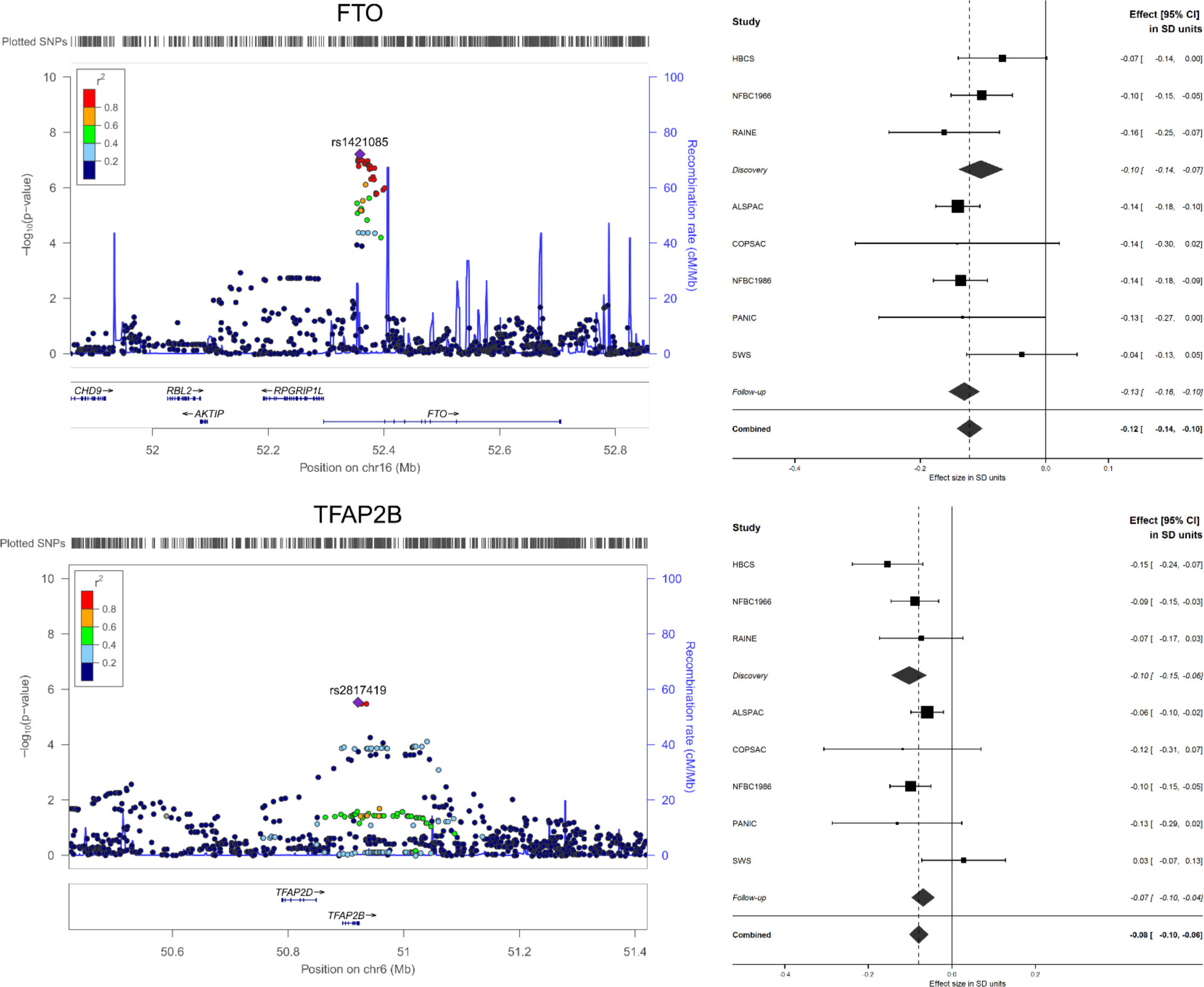
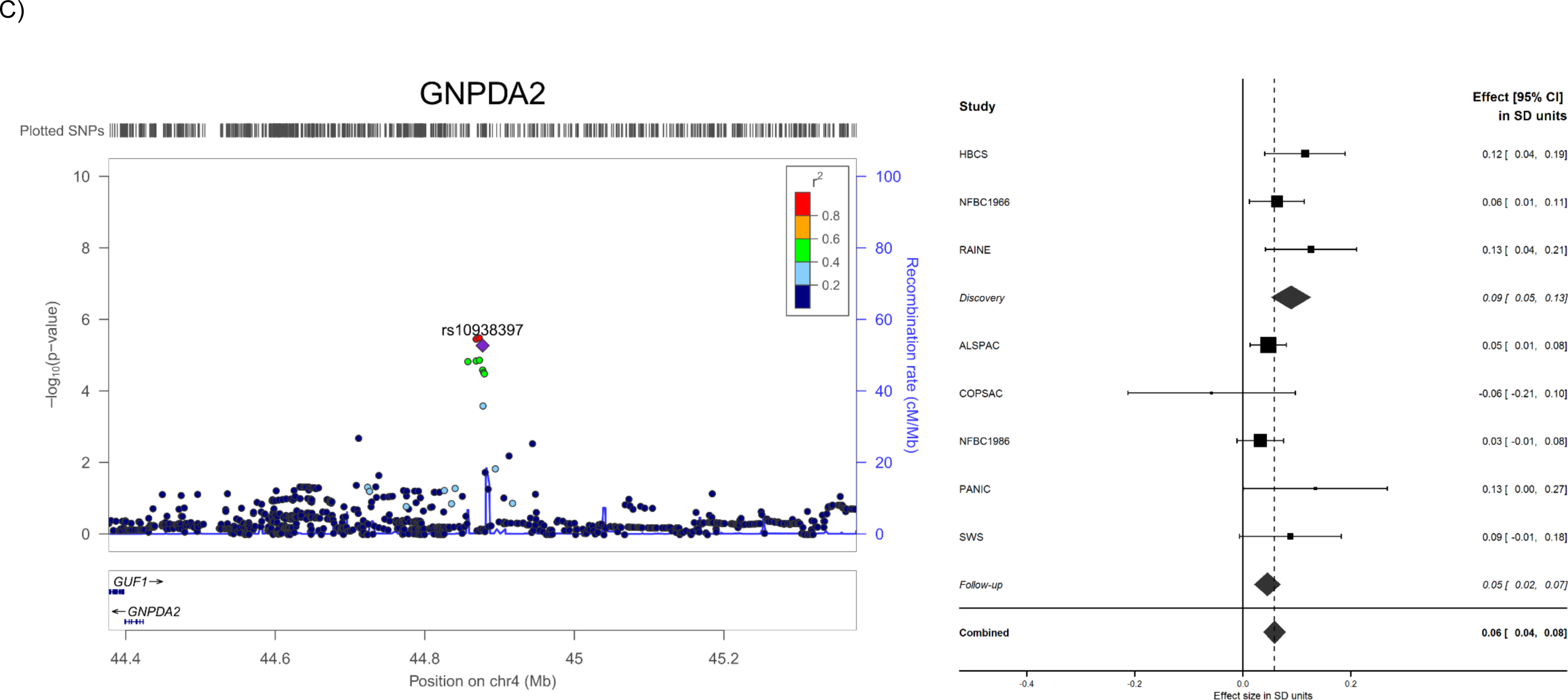
Regional association and forest plots of the four genome-wide significant loci associated with A) BMI at AP^*^, B) AGE at AR and C) BMI at AR. Purple diamond indicates the most significantly associated SNP in stage 1 meta-analysis, and circles represent the other SNPs in the region with colouring from blue to red corresponding to *r* values from 0 to 1 with the index SNP. The SNP position refers to the NCBI build^36^. Estimated recombination rates are from HapMap build 36^78^. Forest plots from the meta analysis for each of the identified loci are plotted abreast. Effect size [95% CI] in each individual study, discovery, follow-up and combined meta-analysis stages are presented from fixed effects models (heterogeneity of the SNP rs9436303 in *LEPR/LEPROT*, see **Supplementary Figure 4**). *At this locus there was heterogeneity between the studies in discovery (I2=72.1%, P=0.01) and combined stage (*I^2^*=59.3%, P=0.002) fixed effect meta-analyses that was mainly due to LISA-D, EDEN and the larger well-defined NFBC1966 study (**Supplementary Figure 4 a & d**). Removing the studies that showed inflated results from meta-analyses did not change the point estimate (**Supplementary Figure 4 c, f, g**). Both fixed and random effect models gave very similar results (**Supplementary Figure 4b & 4e**).

At the *LEPR/LEPROT* locus, the minor G allele of rs9436303 was associated with higher BMI at AP, 0.07SD per allele (95% CI= 0.04, 0.09), *P*_combined_ = 5.1 x 10^−9^, equivalent to a difference of 0.07 kg/m^2^ (95% CI= 0.05, 0.09) per effect allele. The same allele is also associated with earlier age at menarche (*P*= 0.00019 in the ReproGen consortium’s publically available age at menarche GWAS data)^17^ and with increased plasma soluble leptin receptor levels (*P*=1.19 x 10^−9^)^18^ but the SNP was not associated with sex combined adult BMI (*P*=0.96 in the GIANT consortium’s publicly available BMI GWAS data)^1^ amidst a marginal association with BMI in females of <=50 years of age (*P*=0.029) in GIANT. Furthermore, this SNP is in low LD with other common variants at the *LEPR/LEPROT* locus that are also associated with severe early-onset obesity (rs11208659, r^2^=0.008; rs1137100, r^2^=0.128)^19,20^; plasma soluble leptin receptor levels (rs1137100, r^2^=0.128; rs1137101, r^2^=0.151)^18^ and age at menarche (rs10789181, r^2^=0.006)^17^. It is also in low LD with the CRP-associated variant identified in our own previous study (rs12753193, r^2^=0.015)^21^ (**Supplementary Table 4**). Using data from the Northern Finland Birth Cohort (NFBC) 1966 study (N=2585) we also show that the previously published genome-wide significant SNPs associated with other traits in the *LEPR/LEPROT* locus are not associated with BMI at AP, and that our index SNP rs9436303 remains associated with BMI at AP (*P<* 1x10^−6^) after conditioning on those other trait-associated SNPs (**Online Methods and Supplementary Table 5)**. Therefore, to our knowledge, rs9436303 represents a novel, distinct, signal for BMI at AP at this locus.

At the *FTO* locus, the C allele of the index SNP rs1421085 was associated with earlier Age at AR, −0.12 SD per allele (-1.44 months, 95% CI= −1.42, −1.46), *P*_combined_ = 3.12 x 10^−30^, and is directionally consistent with a previous gene-centric study^8^. The *FTO* locus has the largest effect on BMI and obesity risk reported in GWAS carried out to date, and the same allele that is associated with earlier Age at AR is associated with higher childhood and adult BMI^22^. Close proxies for this index SNP have also been associated with adult BMI^23^, type 2 diabetes^24^, metabolic syndrome^25^, waist circumference^26^ and extreme obesity^27^ (**Supplementary Table 4**).

At the *TFAP2B* locus each additional copy of the A allele of our index SNP rs2817419 was associated with earlier Age at AR, −0.08 SD per allele (~-0.96 months, 95% CI= −0.94, −0.98), *P*_combined_ = 4.4 x 10^−11^, and the same allele is robustly associated with higher adult BMI, 0.03 SD per allele (95% CI= 0.02, 0.04), *P*=3.7 x10^−15^ in the GIANT consortium’s BMI GWAS data^1^. The G allele of the adult BMI^1,28^ and waist circumference^29^ associated index SNP rs2207139 in *TFAP2B* from GIANT results in earlier Age at AR in our data, −0.09 per SD (^~^-1.1months, 95% CI: −1.12, 1.08), *P*_discovery_ = 0.0002 (**Supplementary Table 6**). However, rs2207139 is in very low LD (r^2^= 0.035) with our index SNP rs2817491, and these are likely to represent two distinct signals at this locus. In support of this, conditional analyses in the NFBC1986 data adjusting the effect of rs2817419 on Age at AR for rs2207139 and *vice versa* showed that the two SNPs have independent effects on Age at AR (-0.07 SD (95% CI= −0.13, −0.01), *P*=0.03 and −0.10 SD (95% CI= −0.17, −0.03), *P*=0.007 respectively) (**Online Methods**).

Overall, the identified associations at *FTO and TFAP2B* are directionally consistent with observational associations between the timing of AR and adult BMI^9^ (i.e. reaching AR at an earlier age is associated with higher BMI whereas reaching AR later is associated with lower BMI).

At the *GNPDA2* locus, the G allele of the index SNP rs10938397 was associated with higher BMI at AR, 0.06 SD per allele (0.06 kg/m^2^, 95% CI= 0.04, 0.07), *P*_combined_ = 2.9 x 10^−8^, with the same allele being associated with higher adult BMI in previous reports^30^.

There was no heterogeneity between the studies, except at *LEPR/LEPROT* (**Supplementary Figure 3**). For example, in the fixed effect meta-analysis of the discovery stage results for the BMI at AP associated SNP rs9436303 in *LEPR/LEPROT* locus showed heterogeneity (**Supplementary Figure 3a**) and random effect meta-analysis did not change the heterogeneity (**Supplementary Figure 3b**). However, excluding the smaller LISA-D study (N=282) that showed directionally inconsistent effects to the other studies, from the discovery stage meta-analysis for this SNP reduced the heterogeneity but did not change the point estimates (**Supplementary Figure 3c**). Similarly, there was heterogeneity in the combined stage fixed meta-analysis for the same SNP in *LEPR/LEPROT* (**Figure 3d**), and random effect meta-analysis did not change the heterogeneity (**Figure 3e**). Fixed effect meta-analysis excluding the EDEN and NFBC1966 studies that showed inflated results showed no heterogeneity, and the point estimates were similar with and without the two studies (**Supplementary Figure 3f & 3g**). All four SNPs together explain 0.1 %, 0.7% and 0.3 % variance in BMI at AP, Age at AR and BMI at AR, respectively.

### Prioritizing candidate genes and pathways in the four early growth trait associated loci for functional characterization

We first considered the genes that harbor, or are nearest to, the index SNP as a potential candidate. The known biological functions and molecular mechanisms of the four proteins encoded by these are given in **Supplementary Table 7.** However, as the four genome-wide significant signals are located in regions harboring multiple genes with dense linkage disequilibrium (LD) structure, we are unable to locate with certainty the causal gene or gene variant. Therefore, the gene nearest to the lead signals may not necessarily be the gene influenced by the underlying causal variant/s. For example, at the *FTO* locus more than one gene, which includes the *FTO^31,32^, RPGRIP1L^33^ and IRX3/IRX5* in the IRXB gene cluster^34,35^ have been implicated as the candidates for mediating the observed associations with adult adiposity. Given our index SNP at *FTO* associated with Age at AR is highly correlated with the BMI associated SNPs at the same locus, it is plausible that the same genes play a role in the regulation of Age at AR and adult adiposity.

To identify other potential candidate genes and provide further insights into the molecular mechanisms underlying the four genome-wide significant loci associated with BMI at AP, BMI at AR or Age at AR, we searched for association of the index SNP with *cis*–acting expression quantitative trait loci (eQTLs) in five different living tissues: liver, skin, whole blood and subcutaneous and omental fat, and repeated these lookups in 44 post-mortem tissues from the Genotype-Tissue Expression (GTEx) consortium^36^ (**Online Methods**), which mostly reproduced the results of the eQTL association in living tissues. For two out of the four loci analysed, the index SNP was strongly associated (at *P* < 0.001) with the expression of one or more nearby (+/-1 Mb) genes (**Supplementary Table 8**). For example, the BMI at AP associated SNP rs9436303 in the *LEPR/LEPROT* locus was associated with gene expression levels in all five tissues: the strongest association was with *LEPROT* (*P*=1.5 x 10^−51^) gene expression levels in subcutaneous fat. The consistency of the results for this locus across five different tissues and in four different datasets (MuTHER, KORA, deCODE and Kaplan) suggests that these findings are robust and imply that, at least for this locus, the underlying causal variant functions through gene expression. Both the *LEPR* and *LEPROT* genes are good biological candidates for having a role in the regulation of BMI at AP in infancy because the *LEPR* gene encodes the receptor for leptin, an adipocyte-specific hormone that plays a major role in the regulation of appetite and energy balance, reproduction, growth and the immune system^37,38^. Variation in leptin concentrations is also associated with adult body mass^38^ and rare mutations in the *LEPR* gene cause monogenic obesity^39^. The *LEPROT* gene negatively regulates *LEPR* cell-surface expression^40^ and growth hormone (GH) receptor expression in the liver^41^, thereby decreasing hepatic responses to leptin and growth hormone. This is consistent with the finding by Wu *et al*^42^ in mice showing that *LEPROT* could be central to the nutritional regulation of growth and growth hormone binding in the liver and chondrocytes. This evidence further suggests a role for *LEPROT* in the hepatic synthesis of the insulin-like growth factor 1 (IGF-1)^42^. The BMI at AR associated index SNP rs10938397 in *GNPDA2* was associated with *GUF1* (GUF1 Homolog, GTPase) transcription in subcutaneous fat (*P*=7.10 x10^−4^); a gene implicated in mitochondrial protein synthesis (**Supplementary Table 8**).

To explore the biological pathways and networks underlying early growth, we applied a gene set enrichment analysis (MAGENTA)^43^ to the discovery stage GWAS results (**Online Methods**). We identified enrichment of two gene sets (**Supplementary Tables 9 and 10),** suggesting that genes in Age at AR associated regions are involved in the IGF-1 signaling and apolipoprotein pathways (false discovery rate (*FDR*) < 0.05, *P* < 1.6 x 10^−3^) (**Supplementary Figure 6**). The apolipoprotein pathway has an important role in lipid transport and metabolism, while the IGF-1 signaling pathway has a well-established role in neonatal and pubertal growth^44,45^ and in the regulation of energy metabolism through the activation of PI3K/AKT pathway via either the insulin receptor or the IGF-1 receptor^46^. The present finding supporting the overexpression of the IGF-1 pathway with an earlier Age at AR is consistent with the IGF-1 and the high-protein hypothesis exemplified by the work of Rolland-Cachera and colleagues in pediatric nutrition (for a recent review see Rolland-Cachera *et al.* 2016)^47^. This hypothesis suggests that the risk of childhood obesity associated with high-protein intake during lactation could be caused by stimulation of the IGF-1/GH pathway leading to an early age at adiposity rebound. It is therefore possible that higher IGF-1 levels, via genetic and/or nutritional factors, might reduce GH levels and expression via a negative feedback^48,49^. Subsequent, lower circulating levels of GH might also suppress lipolysis and contribute to fat accumulation^50,51^, potentially affecting normal BMI trajectories and Age at AR, and thereby risk of adult obesity and metabolic disorders.

In a look up of the index SNP, and the top 10 intragenic SNPs associated with Age at AR (at P<0.05) in genes from the IGF-1 and apolipoprotein pathways (**Supplementary Note 1 and Supplementary Table 11**) in cis eQTL data (**Online Methods**), a total of 29 eQTLs were associated with 16 different transcripts (all with *P* < 1 x10^−3^). The most significant association was observed between *APOL4* RNA abundance in subcutaneous adipose tissue and rs132700 near the same gene in the apolipoprotein pathway (*P* = 1.02 x 10^−11^) (**Supplementary Table 12**). One SNP within *PTK2* of the IGF-1 signaling pathway was strongly associated with omental and subcutaneous adipose tissue expression of Argonaute 2 (*EIF2C2*), a nearby gene encoding a protein involved in regulation of microRNA transcription. The SNPs within *HDLBP, APOB* and *APOL4* of the apolipoprotein pathway and SNPs in *IGF1R, PTK2, GRB10* in the *IGF-1* signaling pathway are all likely to affect DNAse activity and chromatin state and expression, while SNPs within *HDLBP, APOB* and *APOL4* are also likely to affect transcription factor binding according to RegulomeDB^52^ and Haploreg^53^.

### Association of the adult BMI associated loci with early growth traits

In a look-up analysis of the 97 adult BMI associated loci from the GIANT consortium^1^ in our stage 1 GWAS meta-analysis data, we observed that an excess of adult BMI increasing alleles was associated (*P*<0.05) with increased BMI at AP (12/97 associations, *P_binomial_* =0.003), BMI at AR (19/97 associations, *P*_binomial_ =3.05 x 10^−7^) and earlier Age at AR (26/97 associations, *P*_binomial_ =1.27 x10^−12^) (**Supplementary table 6**). These were directionally consistent with the reported observational associations between the same early growth traits and adult BMI (**Supplementary Table 13**). We also created weighted genetic risk scores using the 97 adult BMI associated SNPs from the GIANT consortium^1^ in the two NFBC studies (**Online Methods**). When comparing the bottom 20% of the children carrying a maximum of ^~^84 weighted BMI increasing alleles (i.e. bottom quintile) to the top 20% of the children carrying a maximum of ^~^111 weighted BMI increasing alleles (i.e. top quintile), there was a mean difference of 0.02 SD (95% CI= 0.01, 0.03) in PWV; 0.16 SD (95% CI= 0.07, 0.24) in Age at AP; 0.14 SD (95% CI= 0.05, 0.22) in BMI at AP; −0.64 SD (95% CI= −0.72, −0.56) in Age at AR and 0.55 SD (95% CI= 0.47, 0.63) in BMI at AR (**Supplementary Figure 4**). Furthermore, we observed correlations (at *P*<0.0001) between the effects of the 97 adult BMI associated SNPs, and Age at AR and BMI at AR (**Figure 4**) but no correlations were observed with the effects on the other early growth phenotypes (**Supplementary Figure 5**). Taken together our results suggest that the variants involved in the regulation of adult BMI have effects that begin in early childhood, around the age at AR.

**Figure 4.**
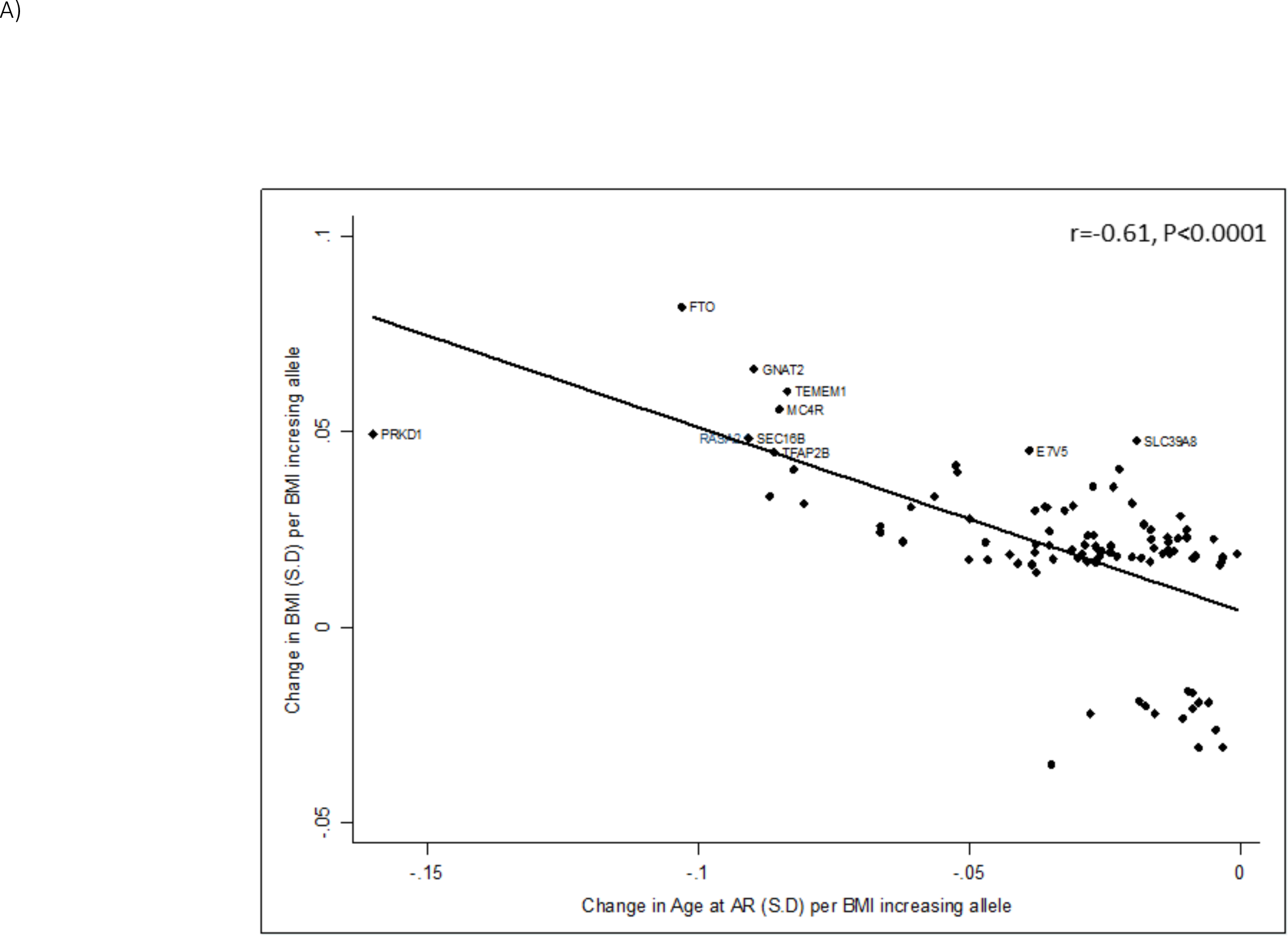
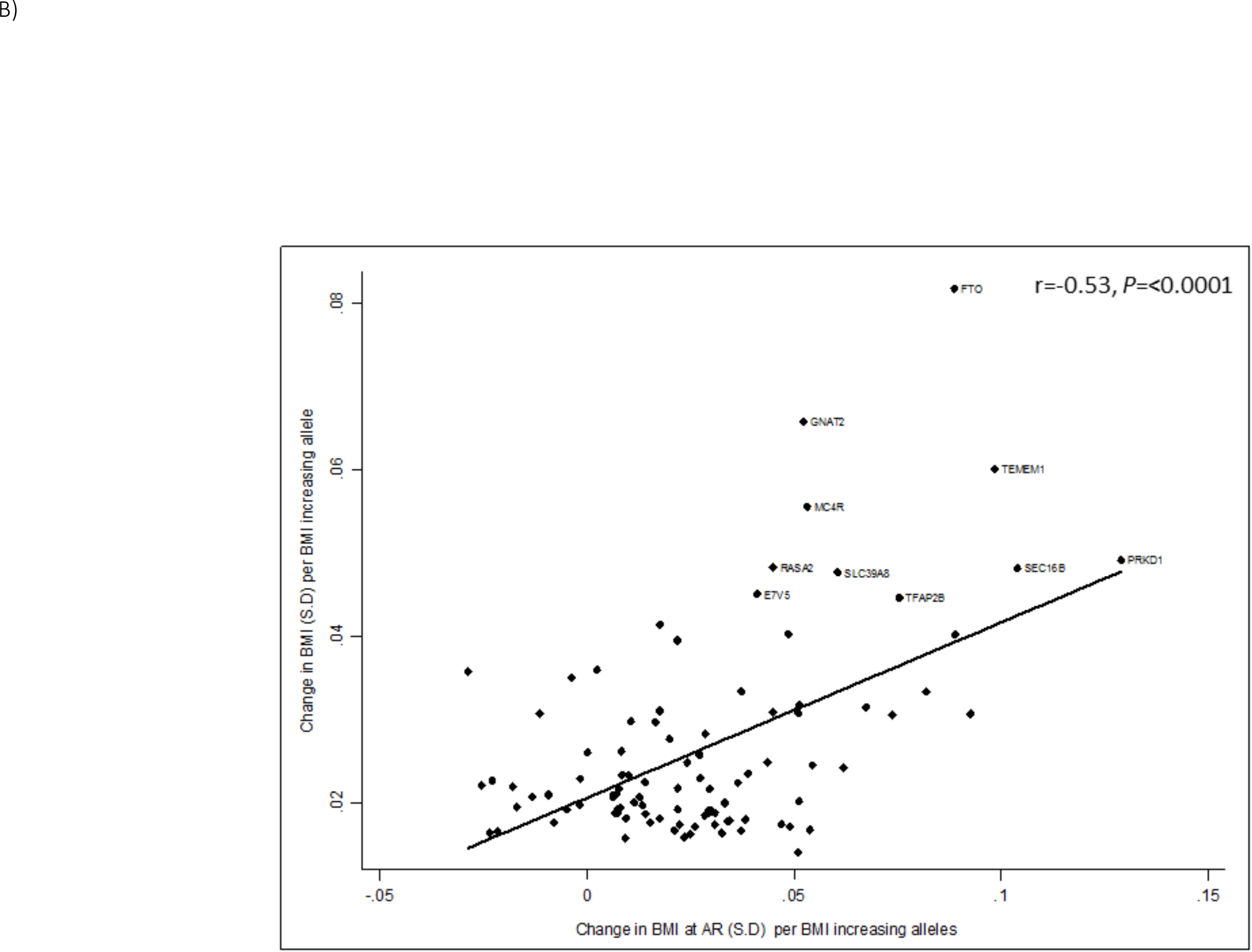
Scatter plots of the effect size estimates (SD units) of the 97 adult BMI associated loci on adult BMI in GIANT consortium^1^ and Age at AR (A) and BMI at AR (B) in the current discovery stage GWAS meta-analyses. The effect size of the adult BMI increasing allele is plotted. The *r* represents the correlation coefficient between the effect size of the early growth phenotype and adult BMI and *P* is P-value for this correlation coefficient. The scatter plots of the other early growth phenotypes are given in **Supplementary Figure 5**.

### Genetic link between early growth and health outcomes

To further explore the genetic links and, thereby, help prioritize the potential causal relationships between early growth and adverse health outcomes, we estimated their genetic correlation using LD score regression (**Online Methods**). In contrast to observational associations, genetic correlations between complex traits can be useful in prioritizing observational associations for subsequent causal analyses, as genetic factors are less likely to be confounded and are not altered by the outcome. Here we summarize the key genetic correlations at 1% FDR, and full results are presented in **Figure 5** and **Supplementary Table 14**. We uncovered several genetic correlations that were directionally consistent with the reported observational associations between the same traits (**Supplementary Note 2 and Supplementary table 13**).

**Figure 5:**
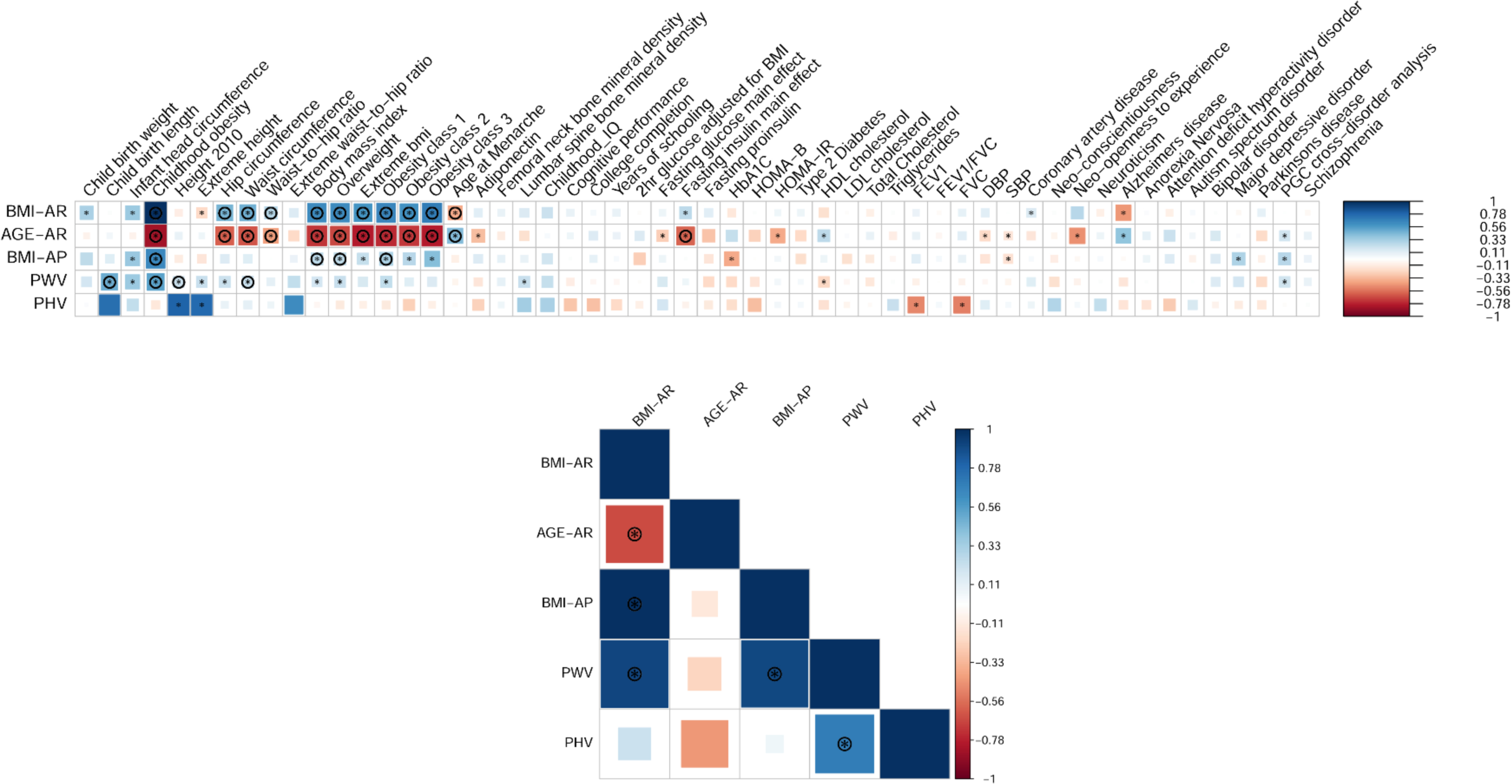
Genetic correlations between five early growth traits and 54 other GWAS traits analyzed by genome-wide association statistics. Blue, positive genetic correlation; red, negative genetic correlation. The correlation matrix underneath represents the genetic correlation among the five early growth traits themselves. The size of the coloured squares is proportional to the *P*-value where larger squares represent a smaller *P*-value. Genetic correlations that are different from 0 at *P*<0.05 is marked with an asterisk. The genetic correlations that are different from 0 at a false discovery rate (FDR) of 1% FDR are marked with a circle. Genetic correlations represented here are presented in tabular format in **Supplementary Table 14**.

In these analyses PHV was not genetically correlated with any of the traits tested at 1% FDR. Conversely, PWV showed a strong positive genetic correlation with childhood obesity (r_g_=0.53, *P*=7.79 x 10^−8^) and was positively correlated with birth length (r_g_=0.50, *P*=0.0003), adult height (r_g_=0.21, *P*=0.0009) and waist circumference (r_g_=0.22, *P*=0.0008). We found very little evidence that PWV was genetically correlated with cardiometabolic traits. The shared genetic contribution of Age at AP with other traits could not be quantified due to low heritability estimates.

BMI at AP also showed positive genetic correlations with childhood obesity (r_g_=0.65, *P*=1.19 x 10^−8^), adult BMI (r_g_=0.26, *P*=0.0002), adult overweight (r_g_=0.26, *P*=0.0007) and adult low-risk obesity class 1 (BMI 30.0-34.9kg/m^2^) (r_g_=0.32, *P*=0.0001).

Age at AR showed a strong positive genetic correlation with age at menarche (r_g_=0.44, *P*=1.90 x 10^−9^), and strong inverse genetic correlations with childhood obesity (r_g_=-0.83, *P*=1.50 x 10^−14^), fasting insulin levels (r_g_=-0.58, *P*=8.69 x 10^−6^) and several adult anthropometric traits including adult BMI (r_g_=-0.72, *P*=3.10 x 10^−18^) and waist circumference (r_g_=-0.62, *P*= 8.40 x 10^−12^).

In contrast to Age at AR, BMI at AR revealed an inverse genetic correlation with age at menarche (r_g_=-0.38, *P*=4.48 x 10^−9^) and positive correlations with childhood obesity (r_g_=1.00 P=6.51 x 10^−25^), adult BMI (r_g_=0.64, *P*=1.60 x 10^−15^) and waist circumference (r_g_=0.48, *P*= 6.07 x 10^−10^). Together the genetic correlations of Age at AR and BMI at AR with age at menarche and BMI are directionally consistent given that earlier age at menarche predicts higher adult BMI^54^ and both are associated with adverse cardiometabolic traits in observational studies^55-57^.

In summary, the results from LD score regression analysis define a robust link between the genetics of early growth and the genetics of later BMI and childhood /adult obesity. In contrast, the LD score regression analysis revealed relatively little evidence for a shared genetic link between early growth traits and lipid metabolism, blood pressure, type 2 diabetes or cardiovascular disease.

## Discussion

We report the first GWAS of six early growth phenotypes derived from longitudinal data. In samples of up to 22,769 term-born children of European descent, we exploited a wealth of repeated weight and height data collected from birth until 13 years. These more refined phenotypes better capture childhood growth patterns relevant to later disease risk than single growth measures in childhood such as BMI and height. The four common variants identified here show consistent effects across several studies despite varying study designs, sample collection methods, and containing data obtained from children born at different geographical regions in Europe demonstrating that our results are robust, and are not affected by any differences between studies. One exception to this is the common variant at *LEPR/LEPROT* where we observed some evidence for heterogeneity. However, excluding the studies that showed inflated results from the meta-analyses reduced or showed no heterogeneity, and did not change the point estimates. Our findings contribute to the understanding of genetic influences on early growth in humans, and provide insights into the underlying molecular mechanisms that link early growth phenotypes with increased risk of obesity in later childhood and adulthood. An exception to this is the common variant in the *LEPR/LEPROT* locus associated with BMI at AP that was not associated with later obesity in childhood or adulthood, although other common variants in this gene are associated with childhood obesity and several metabolic traits, and rare mutations in the same gene cause early onset morbid obesity (OMIM ID: 614963)^39,58^. This raises questions about how the yet-to-be-determined causal variant at *LEPR*/*LEPROT,* or the leptin/free leptin surge in infancy, impact on early development and health. It is noteworthy that research in animals has identified an early peak in leptin concentration, which is thought to be essential not only in regulating energy balance but also in brain development^59^.

Our study has limitations that should be taken in to consideration for future genetic studies of early growth traits. **First**, dense longitudinal growth data meshed with GWAS data are only available in a few cohorts worldwide, so we had limited power to detect genetic variants with smaller effects and/or low allele frequencies. This affected our ability to identify robust associations of some of the promising variants identified in the discovery stage, in particular at *PCSK1* (Table 1). Variants at this locus, which are known to be associated with severe obesity^60^, showed a suggestive association with BMI at AP in our discovery GWAS, but failed to replicate and were not included in further analyses. However, in our discovery stage GWAS for BMI at AP among ~6,222 infants, we had 85% power to detect a variant with an effect size of 0.14 SD at *P*<5 x10^−8^ with minor allele frequency of 0.22. **Second**, it is noteworthy that these derived growth phenotypes are likely to be influenced by a degree of measurement error, especially in cohorts with fewer repeated measures around the time points being estimated, which in turn could have further hampered our power to detect true genetic associations. Despite this, we were still able to discover genetic variants showing robust associations with these derived growth phenotypes. **Third**, in the current GWAS we inferred genotypes based on HapMap Phase 2 imputation; imputation using more comprehensive reference panels can provide greater coverage of the genome and can aid in improving power to detect additional genetic variants. **Fourth**, the present results may not be directly applicable to other ethnic groups as growth patterns and disease risk vary by ethnicity. Trans-ethnic studies may help to identify additional genetic risk variants due to differences in allele frequencies among different ethnic groups. For example, the type 2 diabetes-associated locus *KCNQ1,* which has an effect across multiple ethnic groups, was first discovered in an East Asian GWAS, due to the allele frequency difference between the East Asian and European populations^61^. **Finally**, we did not identify any variants associated with PHV, PWV and Age at AP at genome-wide levels of significance, and this may be due to a combination of smaller genetic effects on growth at this stage of development, reduced statistical power due to smaller sample size or because environmental factors are more influential than genetic influences at this age. The interplay with infant feeding and other environmental factors also warrants additional research.

In conclusion, we have identified three novel loci associated with BMI at AP in infancy at around 9 months, and age and BMI at AR at around 5-6 years, and confirmed the previously reported association of *FTO* with Age at AR^8^, at genome-wide significance levels. The genetic architecture of early growth shares a robust similarity with that of later BMI. For example, the three AR associated loci identified here are strongly associated with adult BMI **(Figure 6**), and the cross-trait genetic correlations based on GWAS data showed the same relationship. The relationship of AR to later adiposity may be due to shared genetic mechanisms or Age at AR and BMI at AR might be on the causal pathway for later adiposity. To test this hypothesis, large independent study populations are required, in which causal analysis methods such as Mendelian randomization can be undertaken using the variants associated with these phenotypes as proxies for early growth phenotypes. However, such studies would currently be underpowered as the variants discovered here only explain a small proportion of the variance in the early growth traits. Taken together, our results suggest that adult obesity has its origins in early childhood, and interventions aiming to promote an optimal growth trajectory in infancy and early childhood could contribute to the prevention of obesity.

**Figure 6.**
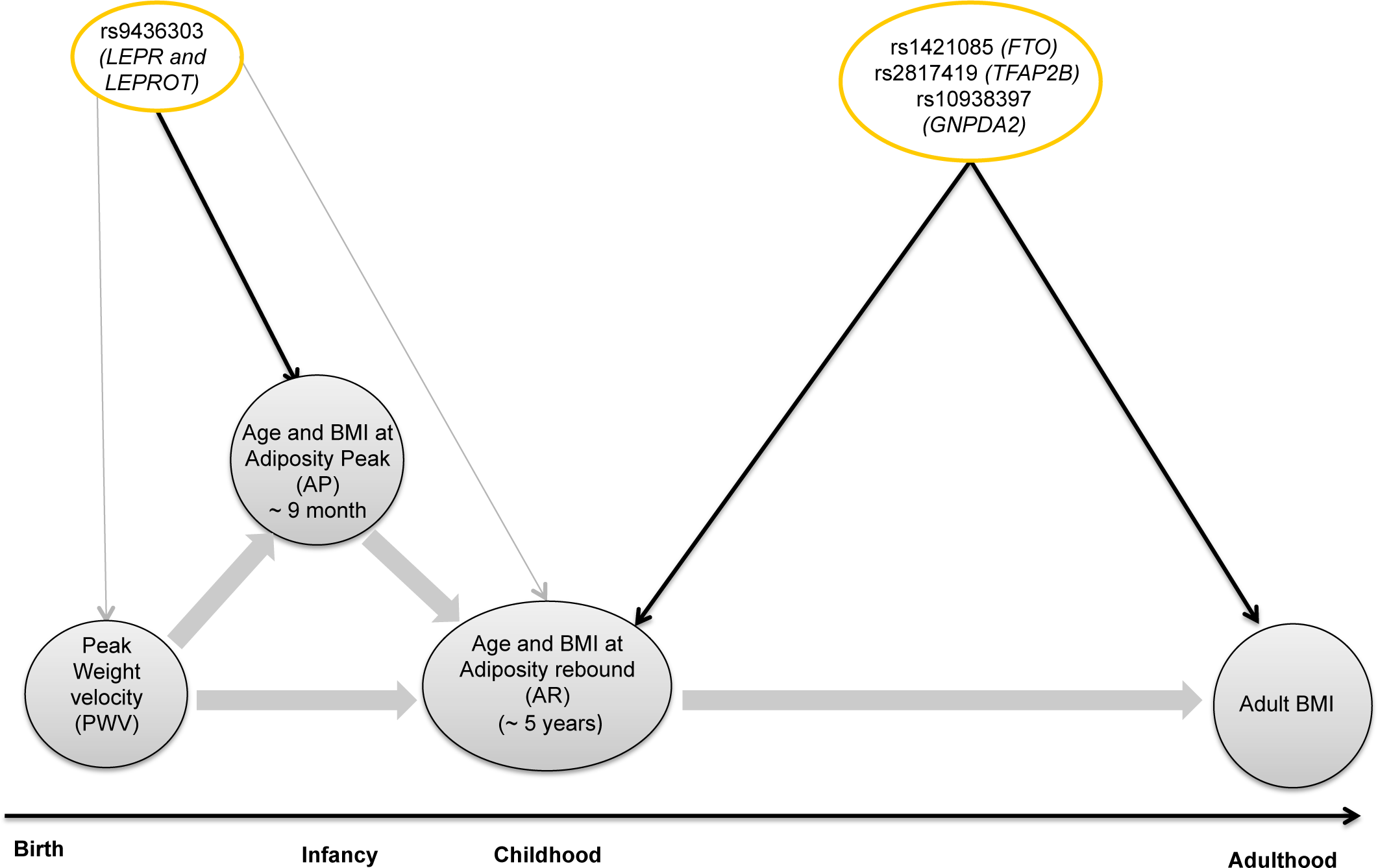
Schematic diagram showing the four genome-wide significant loci associated with early childhood growth phenotypes in the current study and adult BMI in the GIANT consortium. The thick grey arrows show main observational relationships between the early growth phenotypes at *P*<0.0001 in NFBC1966 and NFBC1986 studies, and how early growth phenotypes may be linked with adult BMI. The thin arrows represent the association between the index SNPs and the early growth phenotypes in combined meta-analyses or in look-ups in GIANT consortium data^1^; *P*<0.05 (grey); *P*<0.0001 (black).

## Methods

### Longitudinal growth modeling and derivation of early growth phenotypes

Early growth phenotypes were derived from sex-specific individual growth curves using mixed effects models of height, weight and BMI measurements from birth to 13 years (**Figure 1**). All height and weight data were collected prospectively via either self-reported data or clinical measurements (**Supplementary Table 1 and 2**).These phenotypes were derived separately in each cohort (**Supplementary Note 4)**.

#### Derivation of peak height velocity (PHV) and peak weight velocity (PWV)

The methods for growth modeling and derivation of growth parameters from the fitted curves is described in detail in a previous publication^7^ Parametric Reed1 growth model was fitted in sex stratified non-linear random-effect model as described previously^62^. Term-born singletons (defined as ≥ 37 completed weeks of gestation) with at least three height or weight measurements from birth to 24 months of age were included in the Reed1 model fitting. Maximum-likelihood method for best fitting curves for each individual was used to estimate the growth parameter, PHV (cm/months) and PWV (kg/months).

#### Derivation of age and BMI at adiposity peak (AP) and adiposity rebound (AR)

The methods used for growth modeling of age and BMI has been previously described in detail by Sovio et al, 2011^8^. Due to the specificity of longitudinal changes in BMI *i.e.* succession of peak and nadir as described in figure 1, the data was divided into two age windows for modeling i) growth in infancy using height and weight data from 2 weeks to 18 months of age and ii) growth in childhood using growth and weight data from 18 months to 13 years of age. Each cohort contributed most data available within any of these two age windows. In studies where the data available consisted of both height and weight data within a given window, then the data point nearest to the mid time points of that window were used as a proxy for the BMI measurement. Prior to model fitting, age was centered using the median age of the relevant age window. For example, in the infant growth model at 0-1.5 years, the median age was 0.75 years (which was close to the average age at AP), and in the childhood growth model at >1.5-13 years, the median age was 7.25 years (on average shortly after AR). Linear Mixed Effects (LME) models were then fitted for log-transformed BMI. We used sex and its interaction with age as covariates, with random effects for intercepts *i.e.* baseline BMI, and linear slope *i.e.* linear change in BMI over time. In addition to linear age effect, quadratic and cubic terms for age were included in the fixed effects to account for nonlinearity of BMI change over time.

#### Growth in Infancy

The following model was used to calculate the age and BMI at adiposity peak (AP), and the analysis was restricted to singletons with BMI measures from two weeks to 18 months of age. The model is as follows:

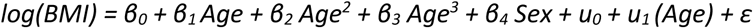

where BMI is expressed in kg/m^2^ and age in year. β_0_, β_1_, β_2_, β_3_, β_4_ are the fixed effects terms, u_0_ and u_1_ are the individual level random effects and ε is the residual error. The age at AP was calculated from the model as the age at maximum BMI between 0.25 and 1.25 year according to preliminary research^7,11^.

#### Growth in Childhood

The model used to measure the age and BMI at adiposity rebound (AR) in childhood is as follows:

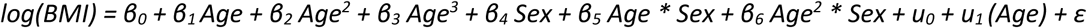

Where BMI is expressed in kg/m^2^ and age in year. β_0_, β_1_, β_2_, β_3_, β_4_, β_5_ and β_6_ are the fixed effects, u_0_ and u_1_ are the individual level random effects and ε is the residual error. Age at AR was calculated as the age at minimum BMI between 2.5 and 8.5 year according to preliminary research^7,11^.

### Stage 1 genome-wide association studies, genotyping and imputation

Stage 1 genome-wide association analyses included up to 7215 children of European descent from five studies (up to four studies for each early growth trait) that had growth data and genome-wide data. These included the Helsinki Birth Cohort Study (HBCS, Finland), Northern Finland Birth Cohort 1966 (NFBC1966, Finland), Lifestyle-Immune System-Allergy Study (LISA, Germany), The Western Australian Pregnancy Cohort Study (Raine, Australia) and Generation R (Netherlands) (**Figure 2**). Informed consent was obtained from all study participants (or parental consent, as appropriate) and the local ethics committees as appropriate approved all study protocols. Study characteristics, genotyping platform, imputation and association test software used, as well as sample and genotyping and imputation quality control steps in each stage 1 study are given in **Supplementary Table 1**. The Stage 1 consisted of a GWAS based on ~2.5 million directly genotyped or imputed SNPs. Imputation of non-genotyped SNPs was undertaken either with MACH or with IMPUTE and were imputed to HapMap Phase 11 CEU reference panel after excluding genotyped SNPs with a minor allele frequency (MAF) < 1%, call rate of at least >95%, and a Hardy-Weinberg Equilibrium (HWE) *P*-value cut off as given in **Supplementary Table 1**.

### Stage 1 genome-wide association analyses and meta-analyses

According to the availability of dense enough data for growth modeling, a total of up to 7215, 6222, 6219 and 6051 children were used to analyse PHV/PWV, Age-AP, BMI-AP and Age-AR /BMI-AR respectively (**Figure 2**). We only included children who were born between 37 and 41 completed weeks of gestation (i.e.: term born) from singleton pregnancies and children who had more than three growth measurements available within the age range in question. Gestational age was either defined from the date of the last menstrual period or ultrasound scans depending on the study. All six early growth traits except for Age-AP and Age-AR were natural log transformed to reduce skewness, and all traits were converted to z-scores prior to association testing to facilitate the comparison of results across the studies. We tested the directly genotyped and imputed variants for association with each of the six early growth traits in a linear regression model assuming an additive genetic effect. The regression models were adjusted for sex and principal components (PC) derived from the genome-wide data to control for potential population substructure (the necessary number of principal components included varied by study). The regression models were also adjusted for gestational age, except for Age-AR and BMI-AR. The genome-wide association analyses (i.e. stage 1) were performed using either SNPTEST or MACH2QTL in each cohort, and data exchange facilities were provided by the AIMS server^63^. All stage 1 study beta estimates and their standard errors were meta-analysed using the inverse-variance fixed effects method in the METAL software^64^. SNPs with poor imputation quality (*e.g.* r^2^< 0.3 for MACH and ‘proper_info’ score < 0.4 for IMPUTE) and/or a HWE *P* <1 x 10^−4^ were excluded prior to the meta-analyses. Double genomic control^65^ was applied: firstly, to adjust the statistics generated within each cohort and secondly, to adjust the overall meta-analysis statistics. Results are reported as a change in standard deviation (SD) units per effect allele as reported in **Table 1**.

### Selection of SNPs for stage 2 follow up

All loci reaching *P* < 1 x 10^−7^ from stage 1 GWAS of each early growth trait were selected for follow-up in stage 2. These included the two SNPs associated with Age-AR in the *FTO locus* (rs1421085) and in the intergenic region between *RANBP3L* and *SLC1A3* (rs2956578), and the SNP associated with BMI-AP in *LEPR*/*LEPROT* (rs9436303). Four further SNPs (one SNP associated with BMI-AP *near PCSK1* (rs10515235), one SNP associated with Age-AR in *TFAP2B* (rs2817419), and two SNPs associated with BMI-AR near *GNPDA2* (rs10938397) and in *DLG2* (rs2055816)) were selected for follow-up on the basis of showing an association with an early growth trait at *P* < 1 x 10^−5^ and being in/near genes with established links to adiposity and metabolic phenotypes except for *DLG2*, a possible candidate gene involved in glucose metabolism^66^. In addition, one locus with a plausible association (*P* = 5.91 x 10^−5^) with PWV, near *TMEM18* (rs2860323), was also selected for follow-up based on previous reports showing an association with severe early onset obesity^19^ and its association with BMI in adulthood^29^ and childhood^6^ (**Supplementary Table 3**). No loci for PHV or AGE-AP passed the p-value threshold or other selection criteria used for follow up. **Supplementary Table 3** shows the SNP selection criteria and proxies used in more detail.

### Stage 2 follow up of lead SNPs in single SNP association analyses

For follow up of lead signals selected from stage 1 we used data from up to 16,550 children of European descent from 12 additional population based studies (up to 11 studies for each early growth trait), namely the Avon Longitudinal Study of Parents and Children (ALSPAC, United Kingdom), Cambridge Baby Growth Study (CBGS, United Kingdom), Children’s Hospital of Philadelphia (CHOP, United States of America), Copenhagen Prospective Study on Children (COPSAC, Denmark), Danish National Birth Cohort (DNBC, Denmark), Étude des Déterminants pré- et postnatals du développement et de la santé de l’ENfant (EDEN, France), The Exeter Family Study of Childhood Health (EFSOCH, United Kingdom), INfancia y Medio Ambiente Project (INMA, Spain), Lifestyle-Immune System–Allergy Study (LISA (R), Germany), Northern Finland Birth Cohort Study 1986 (NFBC1986, Finland), The Physical Activity and Nutrition in Children (PANIC, Finland) and Southampton Women's Survey (SWS, United Kingdom). We used directly genotyped or imputed data for the eight SNPs (or proxies of r^2^ >0.8) selected from stage 1 and tested their association in a total of 5367, 16550, 12256, 12192 children of European ancestry with PWV, BMI-AP, Age-AR, BMI-AR respectively (**Figure 2**). Direct genotyping was performed in some follow-up studies by KBiosciences Ltd. (Hoddesdon, UK) using their own novel system of fluorescence-based competitive allele-specific PCR (KASPar). The call rates for all genotyped SNPs were >95%. Study characteristics, genotyping platform, imputation and association test software used, as well as sample and genotyping and imputation quality control steps in each stage 1 study are given in **Supplementary Table 2**. We used the same methods as in stage 1 for sample selection, genotyping quality control, association testing and meta-analysis.

#### Combined analysis of stage 1 and stage 2 samples

All stage 1 and 2 results were meta-analysed using the inverse-variance fixed effects method in either METAL^64^ or R (version 3.2.0; http://www.r-project.org/). In these combined analyses, loci reaching *P* < 5 x 10^−8^ were considered as genome-wide significant and loci reaching *P* < 5 x 10^−6^ were considered as a suggestive association. Heterogeneity between studies was tested by Cochran's Q tests and the proportion of variance due to heterogeneity was assessed using *I^2^* index for each individual SNP at each stage.

### Estimation of genetic variance explained

The variance explained (VarExp) by each SNP was calculated using the effect allele frequency (*f*) and beta (β) from the meta-analyses using the formula VarExp = β^2^ (1 − *f*) 2*f*.

### Analysis of the association of the index SNP associated with early growth phenotypes in other GWAS data sets

We looked up the index SNP or a proxy associated with each early growth trait in publicly available published meta-analysis data sets to assess traits using the Pheno Scanner available at http://www.phenoscanner.medschl.cam.ac.uk/phenoscanner^67^. We used a *P*-value cut off of <5 x 10^−8^ for displaying the association results and ^r2^ cut off of > 0.6 for proxy SNP lookups from 1000G.

### Conditional analyses

To test whether the index SNP (rs9436303) in *LEPR/LEPROT* locus associated with BMI at AP have an effect on BMI at AP independent of the other trait associated SNPs in the same locus we first, tested the individual association of our index SNP and each of the five previously published genome-wide significant SNPs associated with obesity and metabolic traits in the *LEPR/LEPROT* locus with BMI at AP using a linear regression model adjusted for sex and gestational age in 3459 children from the NFBC1966 study using the same exclusion criteria described above. We next tested whether our index SNP has an independent effect on BMI at AP, by adjusting the linear regression model between the index SNP and BMI at AP for each of the five previously published genome-wide significant SNPs, sex and gestational age.

To test whether our index SNP (rs2817419) and the adult BMI and waist circumference associated index SNP (rs2207139) at the *TFAP2B* locus in GIANT have independent effects on Age at AR we tested the association between rs2817419 and Age at AR in a linear regression model adjusting for rs2207139 and sex and *vice versa* in the NFBC1986 study.

### Expression quantitative locus (e-QTL) analysis

To study the molecular mechanisms underlying the significant genetic variants associated with growth patterns, we searched for cis eQTL using results obtained on liver, skin, whole blood and subcutaneous and omental fat living tissue made available by the MuTHER^68^, KORA^69^, DeCode^70^ and Lee Kaplan^71^ studies. Additional lookups were conducted in 44 post-mortem tissues from GTEx consortium^36^. The association analysis were performed with the index SNP. The analysis of eQTL was limited to genes in cis within a +/- 1Mb window of the index SNP

### Pathway enrichment analysis

To explore the pathways associated with early growth phenotypes, we applied a Meta-Analysis Gene-set Enrichment of variaNT Associations (MAGENTA (version 2))^43^ to the stage 1 GWAS results. Briefly, each gene in the genome is mapped to a single SNP with the lowest *P*-value within a 110 Kb upstream or 40kb downstream window of the gene. The corresponding *P*-value, representing each gene, is corrected for confounding factors such as gene size, LD patterns, SNP density and other genetic factors. The adjusted *P*-values are ranked and the observed number of genes in a given pathway above a specified *P*-value threshold (75^th^ and 95^th^ percentiles used) is calculated. This number is compared with that from repeating the process based on 10000 randomly permuted pathways of identical size. In doing so, an empirical gene set enrichment association (GSEA) *P*-value for each pathway is computed. In our study, individual pathways with a FDR < 0.05 and nominal GSEA *P* < 0.05 were deemed significant, and, unless otherwise stated, results for the 95^th^ percentile cut-off analysis are reported. To test whether SNPs in LD with the index SNP were driving the enrichment of the significant pathways the top 10 intragenic SNPs associated with the relevant early growth phenotype (P<0.05) within genes of the significant MAGENTA pathways were selected by the regression models (**Supplementary Note 1**). These were then examined for association with cis eQTL data in the liver, skin, subcutaneous and omental fat dataset as described above, and a proxy (r^2^>0.5) was used if the top SNP was not available in the expression study. In addition, using all top intragenic SNPs and the index SNPs associated with the significant MAGENTA pathways, we searched the regulomeDB^52^ and Haploreg^53^ database for relevant functional data, including: coding variation, regulatory chromatin marks, DNaseI hypersensitivity, protein binding and motif alteration data.

### The association of BMI genetic risk score with early growth traits

We calculated the weighted genetic risk scores in children from NFBC1966 and NFBC1986 studies separately using the 97 SNPs associated with BMI at genome-wide levels of significance in the GIANT consortium in studies of up to 339,224 individuals^1^. A proxy SNP (r^2^>0.8) was used if the index SNP associated with BMI was not available in the NFBC studies. Only 95/97 were used in the NFBC1986 cohort as no suitable proxies were available for two SNPs (rs6477694 and rs2112347). We used the same exclusion criteria applied in the GWAS analyses described above and calculated the weighted score of the 95/97 SNPs using the formula 1 below to account for varying effect sizes of the SNPs. The SNPs were recoded to reflect the number /dosage of the BMI increasing alleles for that SNP. The weight (*W*) for each locus is the effect size of the BMI increasing allele of each SNP from Locke *et al*^1^.

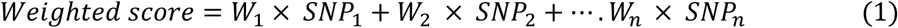

The weighted genetic risk score was rescaled to reflect the number of BMI- increasing alleles using formula 2 as described in Lin *et al^72^*.

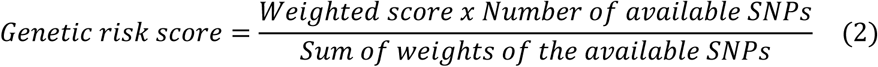

We then created quintiles of weighted genetic risk score in each study containing approximately 20% children in each of the five groups, and tested the mean difference in each early growth phenotype between the lowest and the highest quintile of the weighted genetic risk score in a liner regression model. In the linear regression model we used the quintiles of weighted genetic risk score as the independent variable, and each early growth trait as the dependent variable adjusted for sex and gestational age except for Age-AR and BMI-AR where the regression model was only adjusted for sex. The genetic risk score analyses were carried out in StataMP 13 for Windows (StataCorp, Brownsville, TX). The results from NFBC1966 and NFBC1986 cohort was meta-analysed using the fixed effect inverse variance estimator implemented in the Stata command, “metan”.

### Linkage-disequilibrium (LD) score regression analyses

The use of LD score regression in estimating the genetic correlation between two traits have been described previously^73,74^. Briefly the LD score regression uses GWAS meta-analysis summary statistics of several million SNPs of the two traits under comparison (here the GWAS meta-analysis summary statistics of each early growth trait and the other trait under comparison) and calculates the cross product of the test statistics at each SNP. This cross product of the test statistics is then regressed against the LD score of each SNP (i.e.: sum of the LD r^2^ between a variant and all the variants in a 1 cm region in the genome) and the slope of this regression line gives the genetic correlation between the two traits of interests. We primarily used the LD hub^75^ available at http://ldsc.broadinstitute.org to quantify the genetic correlation between each of the five early growth traits, and health related outcomes. LD hub is a centralised database of summary-level GWAS results from 36 GWAS consortia, and provides a web interface to automate multiple LD score regression analyses in a single run. The analyses can be conducted on selected phenotypes of interest or can be carried out on all traits available on the database in hypothesis free tests. We first reformatted GWAS summary statistics for each of the five early growth traits according to the sample input format provided on the developer’s website prior to uploading on their server. The diseases/traits of interest for LD score regression analyses were selected from the pre complied list of GWAS available on LD hub where summary statistics are available in the required format for running LD score regression. The LD hub uses pre-calculated LD scores based on European samples. We selected a total of 49 disease/traits of interest from 33 GWAS studies in the following pre-compiled categories: education, anthropometric traits, lipids, glycaemic traits, bone mineral density, neurological / psychiatric diseases and other traits (including adiponectin, CAD, T2D, menarche) and submitted the request for LD score regression for each of the five early growth traits at a time. Following each analysis a genetic correlation matrix between the early growth trait and the selected disease/traits were returned which included the genetic correlation value (*r_g_*) its standard error and the corresponding *P*-value for each trait comparison.

However, LD hub currently does not harbour GWAS summary statistic data for lung function measures or blood pressure traits. To quantify the genetic correlation between early growth phenotypes and these measures we obtained GWAS summary statistics data from the SpiroMeta consortium for the lung function measures of FEV_1_, FCV, and FEV_1_/FCV ratio^76^. To generate genetic association statistics for SBP and DBP we carried out GWAS of systolic and diastolic blood pressure (SBP and DBP) using 125,334 subjects from the UK Biobank study^77^ (**Supplementary Note 3**). For these LD score regression analyses we used the Python scripts provided on the developer’s website at https://github.com/bulik/ldsc. Prior to running the LD score regression analyses each summary statistics file was reformatted using the munge_sumstats.py Python script which filtered the SNPs to HAPMAP 3 SNPs as recommended on the developer’s website to minimise any bias from poor imputation quality. SNPs were also excluded if MAF<0.01, ambiguous strand, duplicate rsID and reported sample size is less than 60% of the total available. If the sample size for each SNP was available we used the –N-col to specify the relevant sample size column in the GWAS summary statistics file, and when no sample size column was available we used the maximum sample size reported in the GWAS meta-analysis. After the GWAS summary statistics files were reformatted we then used the ldsc.py Python script to run the LD score regression analyses between each of the six early growth traits, blood pressure and lung function measures. The pre-complied European LD scores calculated from 1000G data available on the developer’s website was used for LD score regression.

## ACKNOWLEDGEMENTS

This publication is the work of the authors, and Marjo-Riitta Jarvelin, Mark McCarthy, Struan Grant, Ken Ong, Vincent Jaddoe and Paul O’Reilly will serve as guarantors for the contents. All the authors acknowledge the following sponsors for their support. The UK Medical Research Council and Wellcome (Grant ref: 102215/2/13/2), NIH grant R01 HD056465, Danish National Research Foundation, the NIH Genes, Environment and Health Initiative [GEI, U01HG004423], NIH GEI [U01HG004438], Lundbeck Foundation [R19-A2059] and the Danish Medical Research Council [09-065592]. French Ministry of Research, Institut National de la Santé et de la Recherche Médicale [INSERM], South West NHS Research and Development, Exeter NHS Research and Development, The Netherlands Organization for Health Research and Development (VIDI 016. 136. 361), the European Union’s Horizon 2020 research and innovation programme under grant agreement No 633595 (DynaHEALTH) and No 733206 (LIFECYCLE); European Research Council (ERC Consolidator Grant, ERC-2014-CoG-648916); Academy of Finland [project grants 209072, 129255 grant] and British Heart Foundation and the Academy of Finland [grants 134839 and 129287], the National Public Health Institute, Helsinki, Finland; Instituto de Salud Carlos III [CB06/02/0041, FIS PI041436, PI081151, PI041705, and PS09/00432, FIS-FEDER 03/1615, 04/1509, 04/1112, 04/1931, 05/1079, 05/1052, 06/1213, 07/0314, and 09/02647], Spanish Ministry of Science and Innovation [SAF2008-00357], European Commission [ENGAGE project and grant agreement HEALTH-F4-2007-201413], Fundació La Marató de TV3, Generalitat de Catalunya-CIRIT 1999SGR 00241. Federal Ministry for Environment [IUF Düsseldorf, FKZ 20462296], Federal Ministry for Environment [IUF Düsseldorf, FKZ 20462296]. National Health and Medical Research Council of Australia [Grant ID 403981 and ID 003209] and the Canadian Institutes of Health Research [Grant ID MOP-82893],, Academy of Finland [project grants 104781, 120315, 129269, 1114194, Center of Excellence in Complex Disease Genetics and SALVE], University Hospital Oulu, Biocenter, University of Oulu, Finland [75617], the European Commission [EURO-BLCS, Framework 5 award QLG1-CT-2000- 01643], NHLBI grant 5R01HL087679-02 through the STAMPEED program [1RL1MH083268-01], NIH/NIMH [5R01MH63706:02], the Medical Research Council, UK [G0500539, G0600705, PrevMetSyn/SALVE] and the Wellcome Trust [project grant GR069224], EU Framework Programme 7 EurHEALTHAgeing 277849. This research has been conducted using the UK Biobank Resource. We thank SpiroMeta consortium for providing the GWAS summary statistics data of lung function measures. Cohort specific acknowledgements are given in the **Supplementary Note 5**.

## AUTHOR CONTRIBUTIONS

All authors contributed in reviewing the paper. **Supplementary Table 15** presents the contribution of each individual author. The names of individuals who contributed to specific statistical analysis and drafting the paper are given below.

***Writing and discussion group on analyses specific to the project***: N.M.G.D.S., S.S., S.D., A.D.S.C.A., U.S., M.K., I.J.M., J.B., R.C., H. R.T., A.M.L., A.R., A.B., V.W.V.J., J.F., P.O., K.O., S.G., M.I.M and M.-R.J.

***GWAS meta-analysis working group***: S.D., H.R.T., U.S. and V.K.

***eQTL and pathway analyses***: A.D.S.C.A.

***LD score regression, BMI SNP look-ups and genetic risk score analyses***: N.M.G.D.S

***GWAS of blood pressure in the UKBiobank***: J.R.P.

## COMPETEING FINANCIAL INTERESTS

The authors declare no competing interest

